# Activity of Antimicrobial Peptides Decreases with Increased Cell Membrane Crossing Free Energy Cost

**DOI:** 10.1101/258863

**Authors:** Rongfeng Zou, Xiaomin Zhu, Yaoquan Tu, Junchen Wu, Markita P. Landry

## Abstract

Antimicrobial peptides (AMPs) are a promising alternative to mitigating bacterial infections in light of increasing bacterial resistance to antibiotics. However, predicting, understanding, and controlling the antibacterial activity of AMPs remains a significant challenge. While peptide intramolecular interactions are known to modulate AMP antimicrobial activity, peptide intermolecular interactions remain elusive in their impact on peptide bioactivity. Herein, we test the relationship between AMP intermolecular interactions and antibacterial efficacy by controlling AMP intermolecular hydrophobic and hydrogen bonding interactions. Molecular dynamics simulations and Gibbs free energy calculations in concert with experimental assays show that increasing intermolecular interactions via inter-peptide aggregation increases the energy cost for the peptide to cross the bacterial cell membrane, which in turn decreases the AMP antibacterial activity. Our findings provide a route for predicting and controlling the antibacterial activity of AMPs against Gramnegative bacteria via reductions of intermolecular AMP interactions.

---

**A**ntimicrobial peptides (AMPs) have received much attention in light of increasing antimicrobial resistance to common small-molecule antibacterial drugs. AMPs exhibit unique modes of action and can be effective against certain antibiotic-resistant bacterial strains. ^1, 2^ AMPs interact with and cross bacterial cell membranes, ^3^ leading to bacterial death. Recent research on AMPs has centred on structure-function relationships, ^4-5^ and studies found that properties of individual peptide, such as hydrophobicity, charge, and amphipathicity, can affect the activities of AMPs. In addition to the intrinsic properties of individual AMPs, inter-molecular interactions between AMPs could also affect antibacterial activity of the resulting peptide formulation. For example, amyloid-β peptide, a natural antibiotic that protects the brain from infection, ^6^ kills bacteria in its monomeric form, but antibacterial activity is lost when high-order peptide oligomeric aggregates are formed. ^7^ Evidence such as amyloid-β loss-of-function upon oligomerization exemplifies the need to consider inter-peptide interactions in the design of AMPs. However, the relationships between the inter-molecular properties of AMPs (eg., self-aggregation) and antibacterial activity remain elusive.

Theoretical studies have recently shown that AMPs have an increased propensity to assume random coil configurations in solution with a low tendency to have a defined structure, when compared to non-AMPs.^8^ Thus, it would appear that AMPs have a higher propensity for nonspecific inter-molecular interactions that could lead to oligomerization and aggregation. As such, we explored the relationship between peptide aggregation propensity and the resulting antibacterial activity of AMPs both theoretically and experimentally. To this end, we chose magainin II (MGN) as our model AMP to study the relationship between self-aggregation and antibacterial activity. MGN II is a naturally-occurring polypeptide that binds to the bacterial membrane and kills bacteria by disrupting membrane integrity. ^9^ Evidence suggests peptide self-aggregation is mainly determined by intermolecular interactions such as hydrogen bonds, electrostatic forces, hydrophobic interactions, and π-π stacking. ^10^ Therefore, fine-tuning the self-aggregation propensity of AMPs requires precise control of these interactions. We chose to test the effect of peptide self-aggregation on MGN II antimicrobial activity by controlling intermolecular interactions between individual MGN II peptide units. Based on our previous work, ^11^ guanine was chosen as an ideal monomer for linking MGN II peptides together via hydrogen bonding and hydrophobic interacttions between peptides. ^12^ This strategy allows us to test MGN II activity without disrupting the native MGN-2 sequence, while promoting inter-peptide aggregation through inter-guanine interactions. To test our hypothesis, we developed a novel strategy in which 1 through 6 guanine units were synthesized into to the N-terminus of the MGN II peptide (**Figure 1b**) to generate MGNs with different self-aggregation propensities based on the different numbers of N-terminal guanine units. We hypothesized that increased peptide self-aggregation propensity decreases the AMP’s antibacterial activity, which can be explained by the increase the energy cost of the peptide crossing the cell membrane (**Figure 1a**). Once a peptide crosses the cell membrane, it must overcome interactions with itself and with other peptides, and these interactions may significantly affect the internalization propensity of peptides and thus their anti-bacterial activities, with a strong tendency to aggregate.

**Figure 1.**
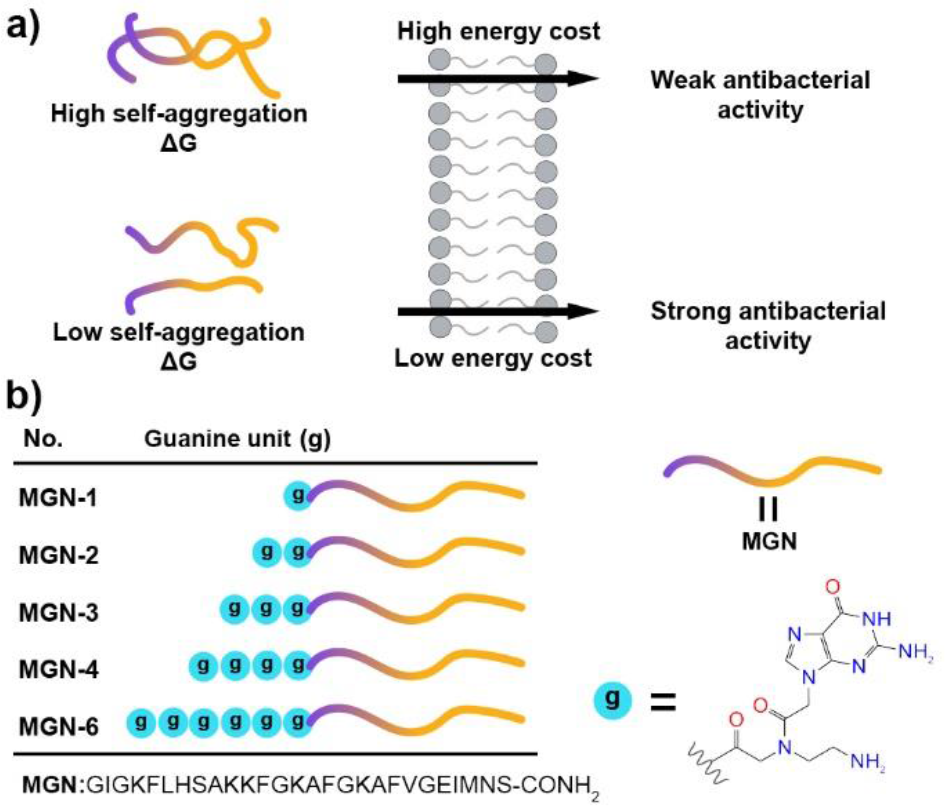
(a) The relationship between self-aggregation of AMPs and their antibacterial activities. (b) Tailor-made peptides were studied as a self-aggregation model. Different numbers of guanine units were attached to the N-terminus as shown. Cyan balls represent guanine units. MGN represents magainin II.

**Figure 2.**
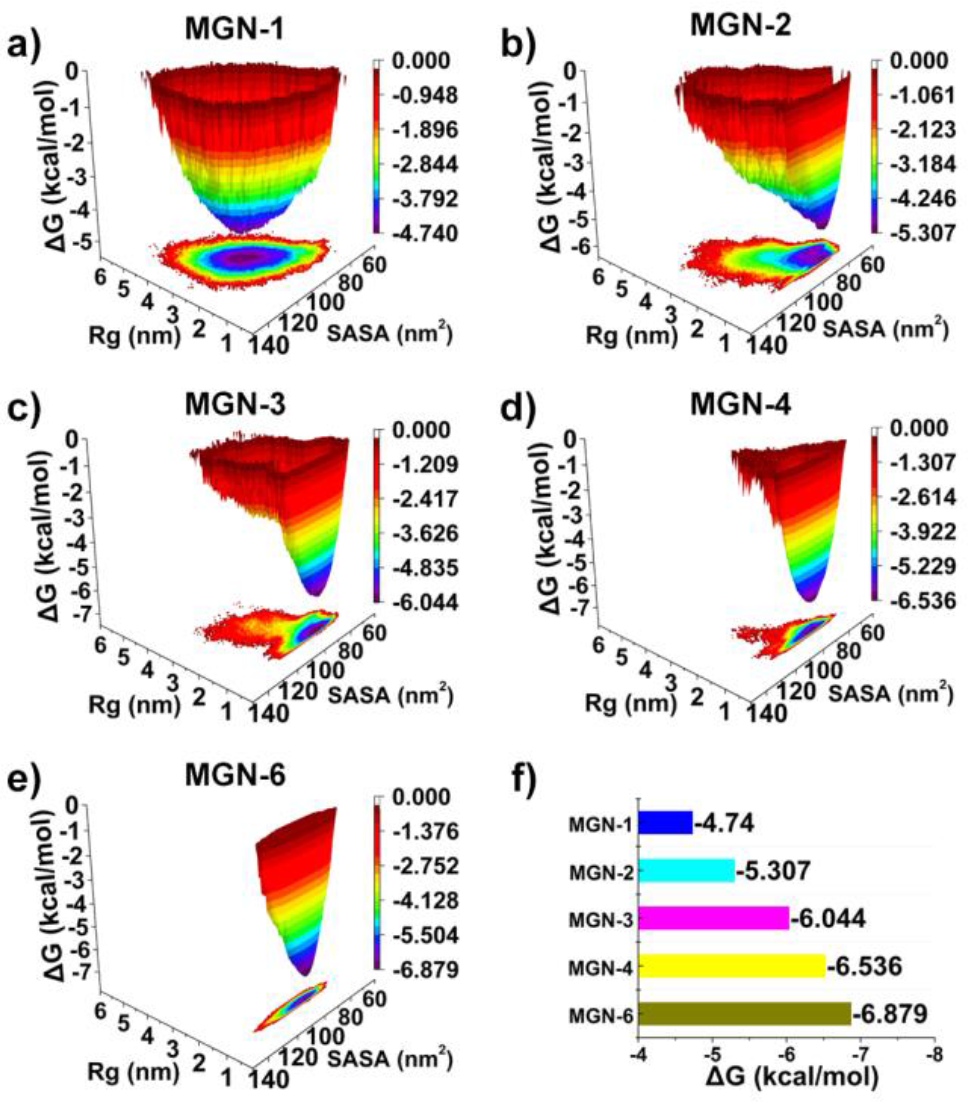
(a-e) Free energy landscape of the MGNs as a function of residue of gyration (Rg) for guanine and solvent-accessible surface area (SASA) of MGNs. (f) Aggregation Gibbs free energy of MGNs.

To test our hypothesis, molecular dynamics (MD) simulations were performed to compare the aggregation Gibbs free energy difference between different aggregated Guanine-tagged antibacterial peptide systems. Comparing the aggregation Gibbs free energy difference between peptides with small sequence differences is normally a very challenging task, thus we implemented a simple but effective strategy inspired by DNA denaturation. ^13^ At higher temperatures, more stable aggregates are less likely to dis-aggregate. For all MGNs, a rapid decrease in solvent-accessible surface area (SASA) indicated the formation of aggregates (**Figure S1a**). However, MGN-1 can quickly disaggregate, as evidenced by the further increase in SASA. Wide fluctuations in SASA usually suggests that the aggregates are not stable, while small SASA fluctuations indicate relatively high inter-peptide stability. ^14^ To visualise the detailed structures of the self-assembled AMPs, cluster analysis was applied to obtain the structure with the highest probability for each system. As shown in **Figure S1b**, guanine units of MGN-1 mostly interact with MGN-II peptide sequence scaffolds. For the other MGN peptides, guanine units interact with each other via hydrogen bonding and hydrophobic interactions. Next, we plotted the free energy landscape of the system using the residue of gyration (Rg) for guanine units and SASA values for the peptides. The free energy landscapes shows that increasing the number of guanine units narrows the free energy wells, indicating that peptide aggregates increase in stability with increasing guanine content. For stable assemblies, the Rg of guanine units adopt a narrow distribution, while the distribution of SASA values is wide, suggesting guanine units form a stable core within the aggregate, while the MGN-II peptide forms a surrounding shell that is relatively flexible. Increasing the number of guanine units decreases the self-assembly Gibbs free energy from -4.740 kcal/mol to -6.879 kcal/mol, indicating that the aggregates become increasingly stable with increasing number of guanine units. As such, our MD simulations establish a quantitative relationship between the aggregation propensity of peptides and the calculated aggregation Gibbs free energy of MGNs.

We synthesized Guanine-tagged peptides to study the MGNs experimentally. Peptides were synthesized by solid phase peptide synthesis (SPPS) ^15^ and purified by high-performance liquid chromatography (HPLC; Table S1–S5; Fig. S2–S6). Circular dichroism (CD) spectra show that the MGNs self-assemble, as evidenced by the negative cotton effect at ~230 nm. ^16^ In the near-UV region (240–320 nm), CD signals mainly reflect guanine-guanine interactions and guanine–MGN-II peptide interactions. ^17^ For MGN-6, guanine units are most likely to form G-quadruplexes due to strong interactions between guanine units. ^18^ Overall, the results of CD experiments are in good agreement with the MD simulations.

Next, antibacterial assays were carried out as described previously for AMPs ^19^ to investigate the antibacterial activities of the Guanine-modified MGNs. MGN-1 displayed the highest antibacterial efficacy, with MIC50 values of 2.2 ng/mL, 17.2 ng/mL, and 42.3 ng/mL against *Escherichia coli, Acinetobacter baumannii*, and *Citrobacter freundii*, respectively (**Table 1**). The MIC50 value of MGN-6 against these three Gram-negative bacteria was higher than 128 negative bacteria was higher than 128 μg/mL in all g/mL in all cases, and activity against *E. coli* was barely detectable, even at the highest concentration, whereas *E. coli* was most sensitive to MGN-1 among the three organisms tested. This phenomenon of decreasing antibacterial activity may be related to the self-assembling propensity of MGNs. Antimicrobial peptide aggregates exhibit less antibacterial efficacy, much like Aβ peptides discussed above lose their antibiotic function in the brain upon aggregation.

**Table 1.** MIC_50_ of MGNs *to Escherichia coli (E. coil), Acineto-bacter baumannii (A.)* and *Citrobacter freundii (C. freundii)*.

| Pathogens | MIC<sub>50</sub> (μg/mL) |  |  |  |  |
| --- | --- | --- | --- | --- | --- |
|  | MGN-1 | MGN-2 | MGN-3 | MGN-4 | MGN-6 |
| *E. coli* | 2.2 | 5.6 | 41.8 | 70.4 | >128 |
| *A. baumannii* | 17.2 | 32.1 | 56.3 | 64.4 | >128 |
| *C. freundii* | 42.3 | 61.8 | 76.4 | 89.5 | >128 |

The antibacterial mechanisms of MGN-1 and MGN-6 were examined using a confocal laser scanning microscope (CLSM) and a LIVE/DEAD BacLight Bacterial Viability Kit. ^20^ The green fluorescent SYTO-9 dye crosses the intact membrane, whereas the red fluorescent propidium iodide enters bacterial cells through lesions in the membrane. As shown in **Figure 3**, both red and green signals were observed in bacteria incubated with MGN-1 at its MIC50 concentration. In contrast, in bacteria incubated with MGN-6 at 128 6 at 128 μg/mL, only green g/mL, only green signal was observed. These results suggest that MGN-1 kills bacteria by crossing the bacterial membrane, but MGN-6 cannot disrupt the cell membrane. These phenomena may be related to the energy cost of peptides aggregates to cross the bacterial cell membrane.

**Figure 3.**
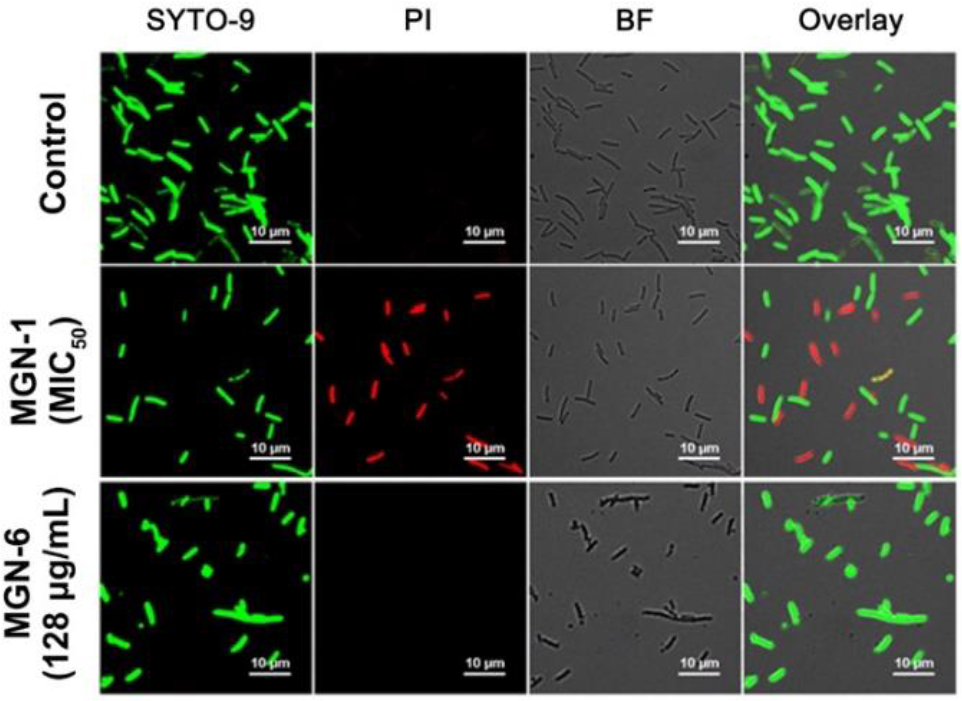
Confocal laser scanning microscope (CLSM) images of *E. coli* cells incubated with 2.2 cells incubated with 2.2 μg/mL MGNg/mL MGN-1 (MIC50 of MGN-1) and 128 cells incubated with 2.2 μg/mL MGNg/mL MGN-6 (the maximum tested concentration of MGN-6) at 37 °C for 1 h and stained with a LIVE/DEAD BacLight Bacterial Viability Kit for 15 min in Mili-Q water. Channel 1 (green), excitation = 488 nm, emission = 500–550 nm; Channel 2 (red), excitation = 561 nm, emission = 570–620 nm.

Next, we determined the energy cost of a peptide crossing the bacterial cell membrane with MD simulations. The permeation Gibbs free energy of a peptide crossing the membrane was calculated using the umbrella sampling method. ^21^ We first carried out a short simulation (50 ns) to probe the interaction between membranes and peptide aggregates, and to obtain a starting structure for further simulation. The peptide aggregates were all attached to the membrane at 50 ns (**Figure 4, Figure S9**). To generate the windows for the umbrella sampling simulation, steered MD (SMD) simulations were carried out with a very slow pulling rate (0.1 nm/ns). The starting structure for the SMD simulation was the final snapshot of the 50 ns unbiased simulation. SMD generated 37 windows, and each window was simulated for 50 ns in the umbrella sampling simulation. The permeation Gibbs free energy increased from 60 kcal/mol to 172 kcal/mol upon increasing the number of attached guanine units from 1 to 6 (**Figure 4, Figure S9**). The peptide must overcome the self-interaction energy with adjacent peptides as it permeates the membrane, and increasing the number of guanine units will increase peptide-peptide interactions, thus the contribution made to the permeation Gibbs free energy increases.

**Figure 4.**
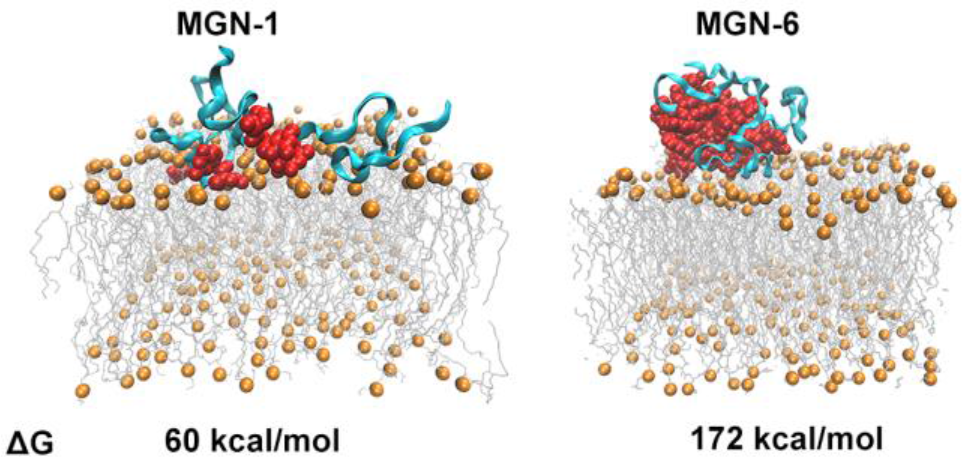
Permeation Gibbs free energy of MGN-1 and MGN-6. Final snapshots of MGNs aggregates attached to the membrane. The MGN II peptide scaffold is colored cyan, guanine units are colored red, and lipids are orange and grey.

Based on our theoretical and experimental results, we find that the antibacterial activity of MGNs is correlated with their aggregation propensity. More precisely, upon increasing the intermolecular interaction propensity of MGNs, the antibacterial efficacy is decreased. To test the breadth of our results, we implemented our experimental strategy to test the anti-microbial activity of another AMP, cecropin A-melittin (CAM). CAM is a hybrid peptide with the sequence KWKLFKKIG-AVLKVL-NH2, which we use to test the validity of our hypothesis on another antimicrobial peptide test case. We synthesized two peptides, CAM-1 and CAM-6 **(Table S6–S7; Figure S10–S11**), with 1 and 6 guanine units as with our MGN tests, and tested CAM-1 and CAM-6 antibacterial activities as described above. We observed that, similar to results obtained with MGN II peptides, CAM-1 also showed greater antimicrobial activity than CAM-6 against several strains of Gram-negative bacteria (**Table 2**). The consistent trend between AMP intermolecular interaction strength and antimicrobial activity for both MGN II and CAM peptides suggests that our understanding of the relationship between intermolecular peptide interactions and antimicrobial activity may be generalizable to other AMPs.

**Table 2.**
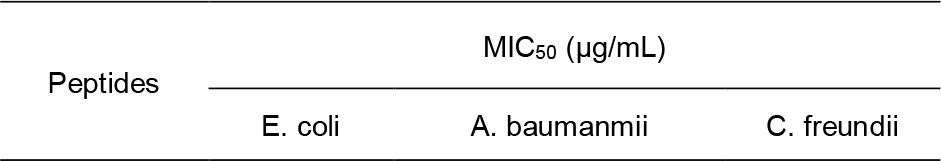
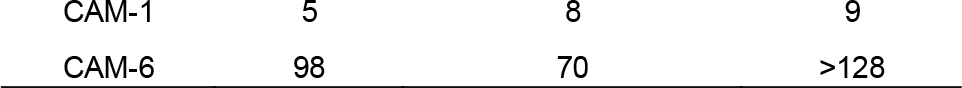
Activity of CAM-1 and CAM-6 against Gram-negative bacteria.

In conclusion, through a combination of theoretical and experimental approaches, we established the relationship between the intermolecular interaction strength and antibacterial activity of AMPs. By introducing different numbers of guanine units, interactions between MGNs can be finely controlled by increasing aggregation propensity, which in turn determines the antibacterial activity. Increasing aggregation between MGNs increases the energy cost of the peptide to cross the bacterial cell membrane, which decreases antibacterial activity. Our method was demonstrated for two unrelated AMP systems. These findings provide a fundamental guiding principle for the design and modification of AMPs.

## Associated content

### Supporting Information

The Supporting Information is available free of charge on the ACS Publications website. Simulation details, experimental details.

## Notes

The authors declare no competing financial interest.

## Acknowledgment

This work was supported by a Burroughs Welcome Fund Career Award at the Scientific Interface (CASI), the Simons Foundation, a BBRF young investigator award, and a Beckman Foundation Young Investigator Award (M. P. L.). M.P.L. is a Chan-Zuckerberg Biohub Investigator. We thank the NSFC (91529101, 21572057 and 21778017) for financial support. R. Z. thanks the China Scholarship Council for financial support. We also thank the Swedish National Infrastructure for Computing (SNIC) for providing computational resources for project SNIC 2016-1-343 and SNIC2016-34-43.

---

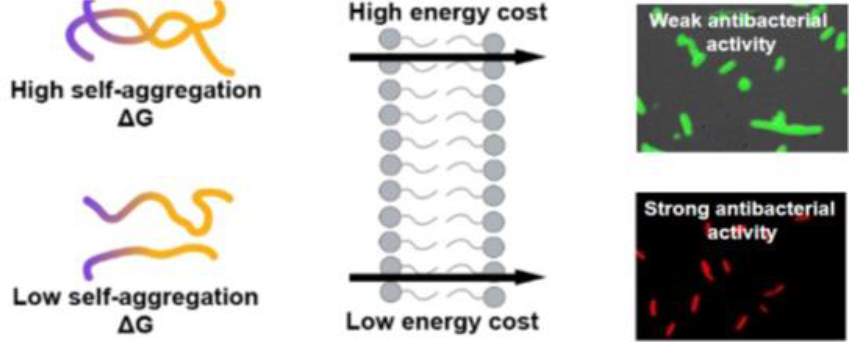

